# Chronic Paraventricular OX1R Overexpression Induces Oxidative Stress and Hypertension in Rats

**DOI:** 10.64898/2026.02.09.704927

**Authors:** Peiying Huang, Nasibeh Yousefzadehkharvanagh, Derrick Simet, Xinqian Chen, Yingli Li, Robert Larson, Lanrong Bi, Qinghui Chen, Bojun Chen, Zhiying Shan

## Abstract

**Background:** Hyperactivity of the orexin system has been implicated in hypertension, yet the long-term impact of orexin 1 receptor (OX1R) overexpression in the paraventricular nucleus (PVN), a key brain region involved in arterial blood pressure (ABP) regulation, remains unclear. This study examined whether chronic OX1R overexpression in the PVN alters cardiovascular, neuroendocrine, and oxidative functions.

**Methods:** Adult male Sprague–Dawley rats received bilateral PVN injections of AAV2-OX1R or control virus AAV2-GFP. Their ABP and heart rate (HR) were monitored for eight weeks via radiotelemetry. Renal sympathetic nerve activity (RSNA) and ABP responses to PVN orexin A administration were assessed using in vivo recordings. Reactive oxygen species (ROS) were quantified using MitoProbe, and gene expression was analyzed by qPCR. Primary neuronal cultures were used to explore underlying mechanism involved in increased ROS production and neuronal activity.

**Results:** PVN OX1R overexpression increased mean ABP (∼10 mmHg), water intake, and ROS levels in both the PVN and peripheral organs. In addition, orexin A evoked exaggerated RSNA (155% vs. 73%) and pressor responses (27 vs. 16 mmHg) in PVN OX1R overexpression rats compared to control rats, these effects were attenuated by the OX1R antagonist SB408124. Plasma vasopressin levels were elevated, and brain sections confirmed increased ROS in OX1R-positive neurons. In vitro experiments further demonstrated that orexin A upregulated ROS-generating enzymes, suppressed antioxidant genes, and enhanced neuronal excitability.

**Conclusion:** Chronic PVN OX1R overexpression induces oxidative stress, sympathetic overactivity, and increased plasma vasopressin, contributing to hypertension development. Targeting PVN OX1R signaling may provide therapeutic benefit for hypertension.

## Introduction

The hypothalamic paraventricular nucleus (PVN) plays a central role in autonomic and neuroendocrine regulation, integrating multiple physiological signals to maintain cardiovascular and fluid homeostasis[1, 2]. Among the neuropeptides involved in PVN function, orexins, specifically orexin A, have emerged as potent modulators of arousal, stress responses, and sympathetic outflow[3, 4]. Orexin exerts its physiological effects through two G-protein-coupled receptors: orexin 1 receptor (OX1R) and orexin 2 receptor (OX2R), with OX1R being highly expressed in key autonomic centers including the PVN[3, 5].

Previous studies have demonstrated that OX1R expression is significantly elevated in the PVN of several models of hypertensive rats such as Dahl Salt sensitive rats and obese Zucker rats[3, 5, 6]. Moreover, microinjection of orexin A into the PVN acutely increases sympathetic nerve activity (SNA) in rats[5, 7]. Conversely, blockade of OX1R with a selective antagonist or genetic knockdown of OX1R in the PVN reduces arterial blood pressure (ABP) and sympathetic outflow[3, 6]. This evidence suggests the potential role of OX1R activation during the development of hypertension. However, it remains unclear whether chronic upregulation of OX1R in the PVN contributes to long-term alterations of cardiovascular function.

It is well established that increased SNA is a major driver of hypertension. In addition, growing evidence indicates that oxidative stress and neuroendocrine disruption also play critical roles in the development of hypertension[8, 9]. Based on this, we hypothesize that sustained activation of orexin signaling may contribute to hypertension through mechanisms involving increased oxidative stress, neuroendocrine imbalance, and impaired regulation of sympathetic nerve system.

Oxidative stress is the harmful state that occurs when levels of free radicals including reactive oxygen species (ROS) exceed the body’s antioxidant capacity. Increased ROS within the PVN is known to facilitate sympathoexcitation and elevate ABP, acting as important mediators in the pathogenesis of neurogenic hypertension[10, 11]. Whether OX1R overexpression can independently drive ROS production and redox imbalance in the PVN, and whether such mechanisms extend to peripheral autonomic system, has not been investigated. Furthermore, orexinergic modulation of neuroendocrine outputs, such as arginine vasopressin (AVP), may amplify fluid retention responses, further contributing to cardiovascular dysfunction[5, 12].

In the present study, we tested the hypothesis that chronic overexpression of OX1R in the PVN induces sustained increases in ABP via enhanced SNA and oxidative stress. Using a combination of telemetry system, recordings of SNA and ABP, microinjection techniques, neuroendocrine assays, oxidative stress imaging, and primary neuronal cultures, we examined the cardiovascular and molecular consequences of chronic OX1R upregulation. Our findings demonstrate that PVN OX1R overexpression elevates ABP, increases ROS production in central and peripheral autonomic system, and alters neuroendocrine responses, thereby implicating orexin signaling as a potential pathogenic contributor to underlying mechanisms of neurogenic hypertension.

## Method

### Animals

Male Sprague–Dawley (SD) rats were purchased from Charles River Laboratories (Wilmington, MA). Animals were housed under a 12 h:12 h light–dark cycle in a climate-controlled room with free access to standard rat chow and water. All procedures followed current NIH guidelines and were approved by the Michigan Technological University Institutional Animal Care and Use Committee (animal protocol number: 00000010))

### Construction and Packaging of AAV2-OX1R Expression Vector

The rat OX1R coding sequence was cloned into an adeno-associated virus (AAV) plasmid driven by the strong, ubiquitous chicken β-actin promoter (CBA) promoter to ensure robust transgene expression. To enable identification of transduced cells, a GFP reporter gene was incorporated into the construct and linked to the OX1R sequence via an internal ribosome entry site (IRES), allowing simultaneous translation of both proteins from a single transcript. This strategy ensured reliable monitoring of expression without interfering with OX1R function. The finalized plasmid constructs were then packaged into AAV2 particles by VectorBuilder, producing a viral vector suitable for targeted in vivo delivery and stable OX1R overexpression in rat brain. A control AAV2-GFP vector, in which the CBA promoter drives GFP expression alone, was used as a control vector.

### Effect of In Vivo AAV-OX1R Delivery into the PVN on ABP and Heart Rate

Eight-week-old male SD rats were anesthetized with inhaled isoflurane (5% induction, 3% maintenance, 1 mL/min in pure O2), and telemetry transmitters were implanted into the femoral artery as previously described[13]. After a 10-day recovery period, rats were randomly assigned to two groups. They were anesthetized and received bilateral PVN injections of either AAV-OX1R or AAV-GFP viral vector (∼10^9^ genomic copy per side). Injections were guided by stereotaxic coordinates: 1.6–1.8 mm caudal to the bregma, 0.2–0.3 mm lateral to the midline, and 7.6–7.8 mm ventral to the dura[5]. BP and heart rate (HR) of each rat were continuously monitored via telemetry for 4 hours one day before viral injection (baseline) and subsequently at the same time slot each week. Eight weeks post-injection, rats were transferred to metabolic cages. After a 24-hour acclimation period, their 24-hour water intake and urine excretion were recorded. The rats were deeply anesthetized with 5% isoflurane in 100% oxygen (flow rate: 1 mL/min) and subsequently euthanized by decapitation. Brain tissue was collected, and PVN samples were punched out for real-time PCR analysis of gene expression. Plasma was also harvested for measurement of AVP and norepinephrine (NE) levels by ELISA.

### Measurement of mRNA Levels by Real-Time PCR

mRNA expression of genes of interest from PVN tissues and cultured neurons were measured by real-time PCR as described previously[5]. Briefly, total RNA was extracted using the RNeasy Mini Kit (Qiagen, Valencia, CA, USA) and reverse-transcribed into cDNA with the iScript™ cDNA Synthesis Kit (Bio-Rad, Hercules, CA, USA). Quantitative PCR was performed on a StepOnePlus Real-Time PCR System (Applied Biosystems, USA) using TaqMan primers, probes, and master mix (Applied Biosystems, USA). All cDNA samples were analyzed in triplicate, and mRNA levels were normalized to GAPDH expression.

### Intracerebroventricular (ICV) Injections and mtROS Measurement

A mitochondrial-targeted fluorescent probes (MitoProbe) synthesized by our research team (Patent No. WO2014063033A2), were used to evaluate the effect of PVN OX1R overexpression on mitochondrial ROS production. MitoProbe emits fluorescence upon binding to ROS, allowing quantification of ROS levels through fluorescence intensity. Eight weeks after viral vector injection (AAV2-OX1R or AAV2-GFP), rats received ICV injection of MitoProbe. Briefly, rats were anesthetized with inhaled isoflurane (5% induction, 3% maintenance, 1 mL/min in pure O2). Then, 10 µL (0.2 µM) of MitoProbe was injected into the right lateral ventricle. Injections were guided by stereotaxic coordinates: 0.8–0.9 mm posterior to bregma, 1.4–1.8 mm lateral to the midline, and 3.2–3.8 mm ventral to the dura, and Mitoprobe were administered at a flow rate of 1 µL/min using an UltraMicroPump3 (World Precision Instruments). Three hours post-injection, the rats were deeply anesthetized with 5% isoflurane in 100% oxygen (flow rate: 1 mL/min) and subseqnently euthanized by transcardial perfusion with 4% paraformaldehyde (PFA). Then brains, adrenal glands, and kidneys were harvested and sectioned for analysis ROS production. Fluorescent images were captured using a confocal microscope Olympus FV1000.

### Preparation of Primary Neuronal Cultures from Neonatal SD Rat Brains

Primary neuronal cultures were isolated from the cerebral cortex of one-day-old SD rats as described previously[14]. Briefly, neonatal pups were deeply anesthetized with 5% isoflurane in 100% oxygen (flow rate: 1 mL/min) and then euthanized by decapitation. their brains were rapidly removed, and cortical neurons were dissociated using the Papain Dissociation System (Worthington Biochemical, USA) according to the manufacturer’s instructions. After isolation, cells were plating in 24-well plate (∼10^5^ cells/well) and maintained in neurobasal medium supplemented with 2% B27 serum at 37 °C in 5% CO₂ incubator for five days before being used for AAV2-OX1R or control vector AAV2-GFP infection. Half of the culture medium was replaced every three days.

### Effect of OX1R Overexpression on Oxidative Stress-Related Genes and Neuroexcitation Markers in Brain Neurons

Five-day-old neuronal cultures were divided into two groups (n=6 per group). One group was incubated with AAV-OX1R (6 × 10 genomic copies/well), and the other group was incubated with an equal amount of control virus (AAV-GFP). Five days after viral infection, each group was further divided into two subgroups: one subgroup received orexin A (100 nM) treatment, while the other received same volume of vehicle control (saline). Six hours after orexin A or vehicle treatment, cells were collected, RNA was isolated, and real-time PCR was performed to assess the expression of genes of interest.

### Renal Sympathetic Nerve Activity (RSNA) Recording

The procedure followed our previously published protocol[5]. In brief, rats were anesthetized with an intraperitoneal injection of α-chloralose (80 mg/kg) and urethane (800 mg/kg). An arterial catheter was inserted into the aorta via the femoral artery and connected to a pressure transducer to measure ABP. HR was derived from the R wave of the ECG. A second catheter was placed in the left femoral vein for drug administration. After tracheal cannulation, rats were paralyzed with gallamine triethiodide (25 mg·kg⁻¹·h⁻¹, IV) and ventilated with oxygen-enriched room air. Anesthetic depth post-paralysis was confirmed by the absence of pressor responses to a noxious foot pinch. Supplemental doses of anesthesia (10% of the initial dose) were administered as needed. End-tidal PCO₂ was maintained within 35–40 mmHg by adjusting ventilation rate (80–100 breaths/min) and tidal volume (2.0–3.0 ml). Body temperature was kept at 37°C using a water-circulating heating pad.

To record sympathetic nerve activity, a left flank incision was made to isolate the left renal sympathetic nerve bundle, which was mounted on a stainless-steel wire electrode (OD: 0.127 mm, A-M Systems) and insulated with silicone-based impression material (Coltene, Light Body). The signal was amplified (P511, Grass Technologies), bandpass-filtered (100–1,000 Hz), notch-filtered (60 Hz), rectified, and integrated (10-ms time constant). Data were digitized at 5,000 Hz using a 1401 Micro3 converter and Spike2 software (v7.04, Cambridge Electronic Design, UK). Background noise was measured at the end of the experiment using a bolus injection of hexamethonium (30 mg/kg, IV), a ganglionic blocker, and subtracted from all integrated values of sympathetic nerve activity. After the experiment, rats were euthanized with an intravenous bolus injection of potassium-chloride (75mg/kg).

### Immunofluorescence Detection of OX1R In the Brain and Adrenal Glands

Immunoreactivity of OX1R in brain and adrenal gland was performed as described previously[5]. Briefly, rats were deeply anesthetized with The rats were deeply anesthetized with 5% isoflurane in 100% oxygen (flow rate: 1 mL/min) and then euthanized by transcardially perfused with 4% PFA in 1× PBS. Brains and adrenal glands were removed and post-fixed in 4% PFA overnight at 4 °C, then transferred to 30% sucrose in PBS at 4 °C until the tissue sank.

Coronal brain sections (20 µm) containing the PVN or Rostral Ventrolateral Medulla (RVLM), as well as adrenal gland cross sections including cortex and medulla, were cut using a cryostat. Sections were washed three times in PBS (10 min each) and incubated for 48 h at 4 °C with rabbit anti-OX1R antibody (Alomone Laboratories, 1:500) in PBS containing 0.5% Triton X-100 and 5% horse serum. After three additional PBS washes, sections were incubated overnight at 4 °C with Alexa Fluor 488 donkey anti-rabbit IgG. Immunoreactivity was examined using a confocal microscope, and micrographs were captured.

### ELISA measurement of plasma AVP and NE

Eight weeks after AAV2-OX1R or control virus injection, rats were deeply anesthetized with 5% isoflurane in 100% oxygen (flow rate: 1 mL/min) and then euthanized by decapitation for trunk blood collection. Blood samples were immediately placed on ice and centrifuged at 2,000 rpm for 15 minutes at 4°C using an Eppendorf 5801 R centrifuge. Plasma was carefully collected and stored at −80°C for subsequent analysis. Plasma levels of AVP and corticosterone were measured using ELISA kits (Thermo Fisher Scientific, USA), following the manufacturer’s instructions.

### Statistical Analysis

Statistical analyses were performed using Prism 6 (GraphPad). Data are presented as mean ± SEM. Two-way ANOVA was used for MAP and HR analysis. Student’s t-tests were applied to evaluate water intake, body weight, gene expression, immunoreactivity, and fluorescence intensity. Differences in MAP and HR for each animal were determined by subtracting baseline values. Fluorescent area quantification was carried out using ImageJ software (version 1.53a). A p-value < 0.05 was considered statistically significant.

## Results

### 1. PVN OX1R Overexpression Leads to Significant Increases in ABP and Water Intake

Adult male SD rats received either an AAV2-OX1R vector or a control AAV2-GFP vector targeted to the PVN. ABP and heart rate (HR) of each rat was measured weekly. The results showed that chronic overexpression of OX1R in the PVN resulted in a mild but statistically significant increase in systolic, diastolic, and mean arterial pressure (MAP) compared with control rats. This MAP elevation peaked at approximately two weeks post-injection (MAP: AAV-GFP: 99.5 ± 0.9 mmHg, n=7, vs. AAV2-OX1R: 107.9 ± 1.2 mmHg, n=7∼10, P<0.05) and persisted through week eight, when the study was terminated. HR remained largely unchanged, except for a transient difference observed in week 6 (Figure 1A).

**Figure 1.**
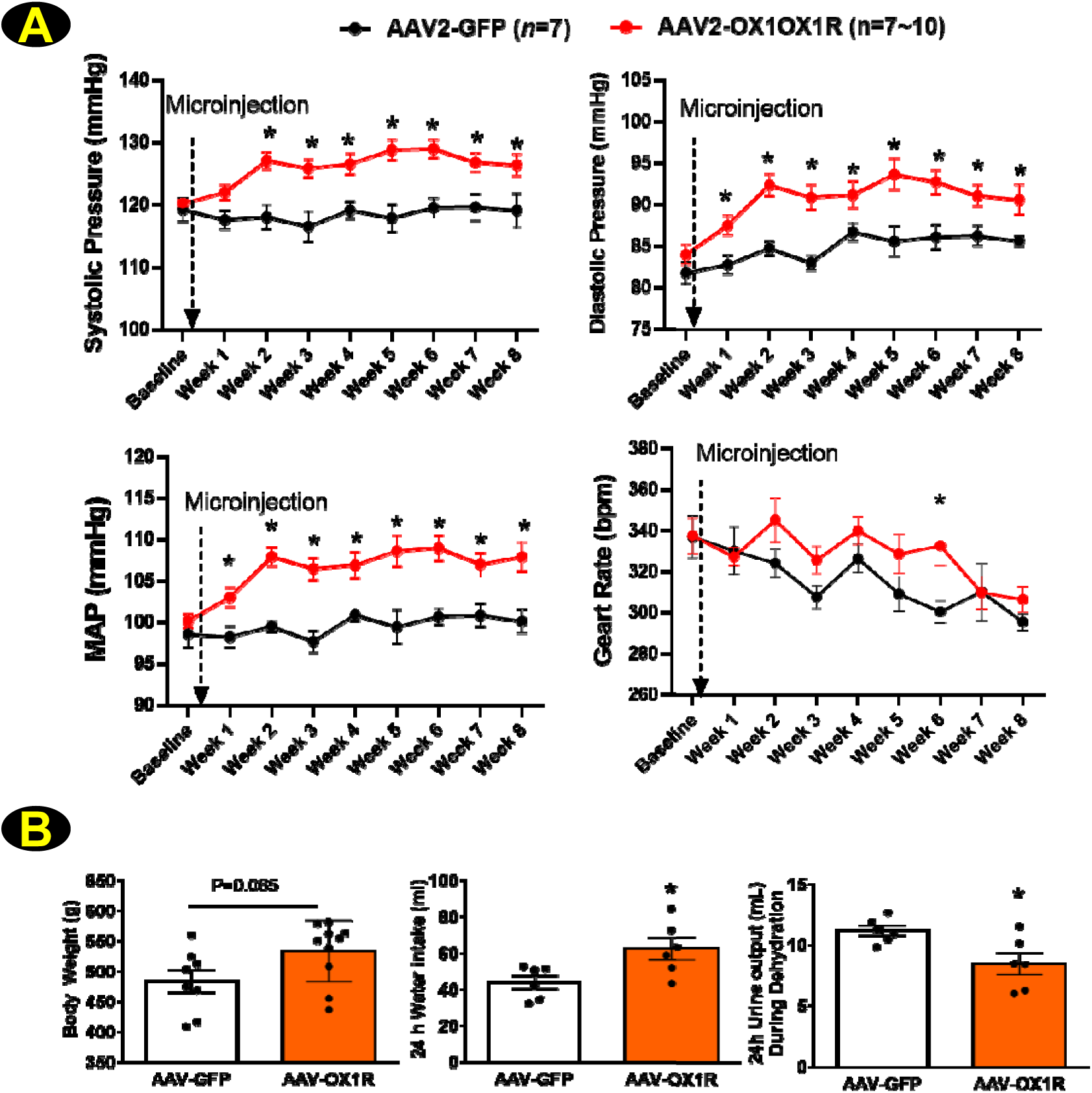
OX1R overexpression in the PVN elevates arterial blood pressure and alters fluid balance in male SD rats. **(A)** Eight-week-old male Sprague-Dawley (SD) rats were randomized into two groups and received bilateral PVN injections of either AAV-OX1R (n = 7–10) or AAV-GFP (n = 7). Arterial blood pressure (ABP) and heart rate (HR) were continuously monitored for eight weeks using a telemetry transducer system. No significant differences were observed at baseline between groups. Beginning two weeks post-injection, the AAV-OX1R group displayed a significant increase in systolic, diastolic, and mean arterial pressure (MAP) compared with controls, peaking at week 3 and remaining elevated through week 8. Data are presented as mean ±SEM, and statistical analysis were performed using two-way ANOVA (*P<0.05). **(B)** PVN OX1R overexpression increased water intake and reduced urine output during water-deprivation testing compared with controls (n = 6 per group, *P < 0.05). In addition, OX1R-overexpressing rats showed a trend towards higher body weight eight weeks after viral vector injection (AAV-OX1R: n = 10 vs. AAV-GFP: n = 8; P = 0.062). Data are presented as mean ± SEM, and statistical analyses were performed using Student’s t-test.

After eight weeks of ABP monitoring, rats were placed in metabolic cages. Following a 24-hour acclimation period, their water intake and urine output were measured. The water intake was significantly higher in PVN OX1R-overexpressing rats compared with controls (AAV-GFP: 44.1 ± 3.7 ml vs. AAV-OX1R: 62.5 ± 6.1 ml; n = 6 per group, P < 0.05). There were no significant changes in urine output when water is free accessible (data not shown). However, during 24 h water deprivation, urine output was significantly lower in PVN OX1R-overexpressing rats (AAV-GFP: 11.2 ± 0.42 ml vs. OX1R: 8.5 ± 0.9 ml; n = 6, P < 0.05) (Figure 1B). Body weight was measured prior to viral vector injection as a baseline and showed no difference between the AAV-OX1R and AAV-GFP groups (data not shown). Eight weeks after viral vector injection, body weight tended to be higher in PVN OX1R-overexpressing rats (AAV-GFP: 481 ± 18.4 g vs.AAV- OX1R: 533 ± 16.0 g; n = 8–10 per group, P=0.065) (Figure 1B), although this difference did not reach statistical significance.

### 2. Verification of PVN-Targeted OX1R Overexpression and Its Impact on Gene Expression

To confirm accurate PVN targeting and successful OX1R overexpression, we evaluated reporter gene (GFP) expression and OX1R mRNA levels. GFP fluorescence verified correct injection site localization, while real-time PCR revealed a 17-fold increase in OX1R mRNA expression in the PVN of AAV2-OX1R rats compared to controls (P < 0.05) (Figure 2A). These results demonstrate successful AAV-OX1R delivery and robust OX1R overexpression in the intended brain region.

**Figure 2.**
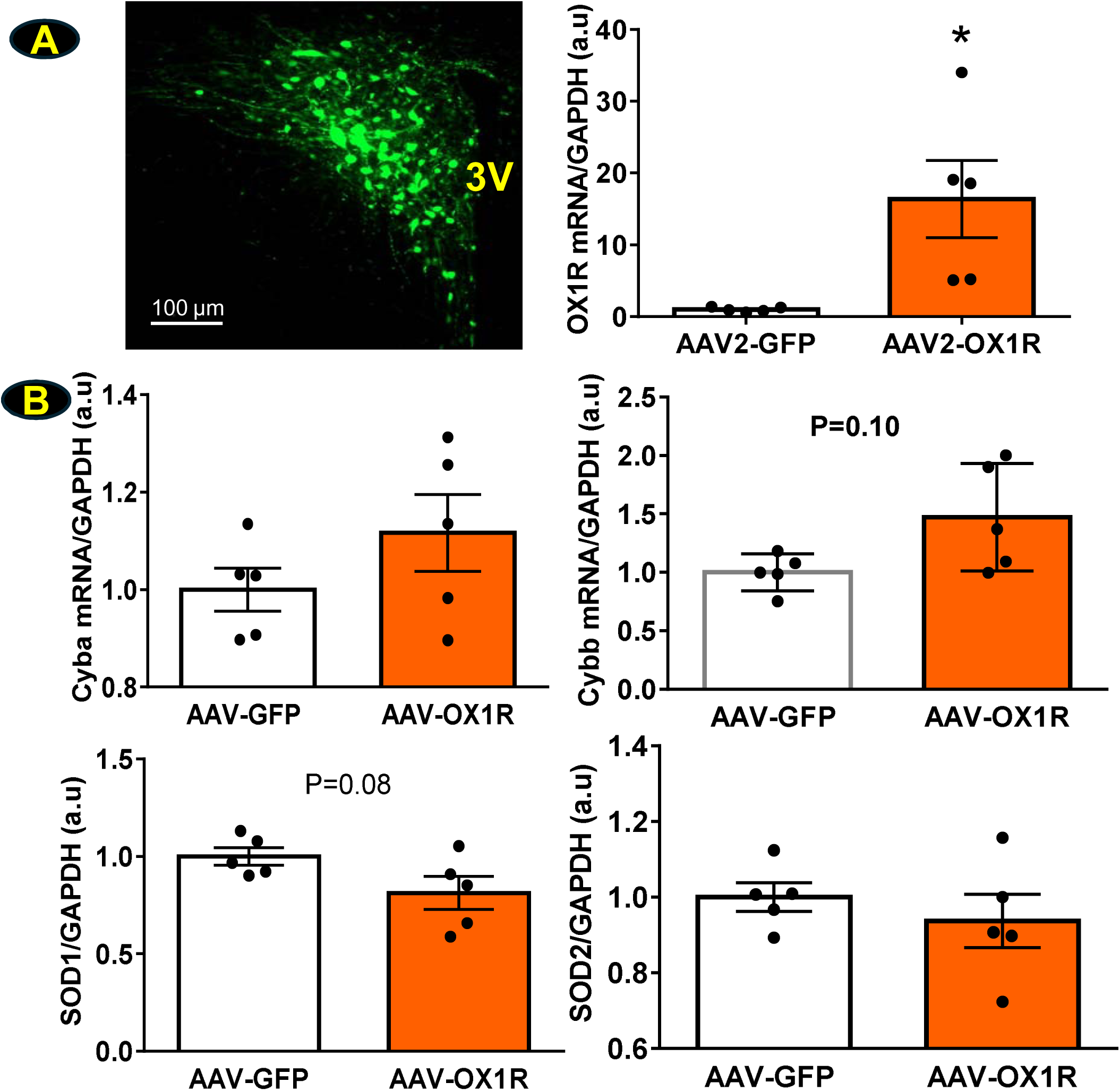
Verification of PVN targeting and OX1R overexpression, and effects on oxidative stress–related genes and neuronal excitation markers in the PVN. **(A)** GFP fluorescence confirmed accurate viral injection within the PVN (scale bar: 100µm). Real-time PCR showed a ∼17-fold increase in OX1R mRNA expression in AAV-OX1R–injected rats compared with AAV-GFP controls (n=5 per group). **(B)** Expression levels of Cyba, Cybb, SOD1, SOD2, and FOS were measured in the PVN. Cybb expression increased 1.6-fold (P = 0.10), and SOD1 expression decreased by 25% (P = 0.08) (n=5 per group). Data are presented as mean ± SEM, and statistical analyses were performed using Student’s t-test.

We further examined whether OX1R overexpression in the PVN influenced genes related to oxidative stress (Cyba, Cybb, SOD1, SOD2). No statistically significant differences were observed. However, Cybb showed a 1.6-fold increase (P = 0.10), and SOD1 expression decreased by 20% (P = 0.08), suggesting a potential role for oxidative stress in OX1R-mediated ABP elevation (Figure 2B).

### 3. PVN OX1R Overexpression Increases ROS Production in the PVN

To further examine the relationship between OX1R expression and oxidative stress, we performed ICV injections of MitoProbe, a fluorescent indicator of reactive oxygen species (ROS), in OX1R-overexpressing rats and AAV-GFP controls. Confocal imaging and quantitative analysis were used to assess ROS levels in the PVN. As shown in Figure 3A–B, AAV-OX1R rats exhibited a significant increase in total ROS fluorescence compared with controls (*P < 0.05). Moreover, ROS co-localization within GFP⁺ cells was approximately ten-fold higher in AAV-OX1R rats (**P < 0.0001, Figure 3C). In the AAV-OX1R vector, OX1R and GFP are co-expressed under a single promoter and separated by an IRES sequence, ensuring that GFP reliably identifies OX1R-overexpressing cells. Collectively, these results indicate that OX1R overexpression induces substantial ROS accumulation in the PVN, particularly within OX1R-expressing neurons

**Figure 3.**
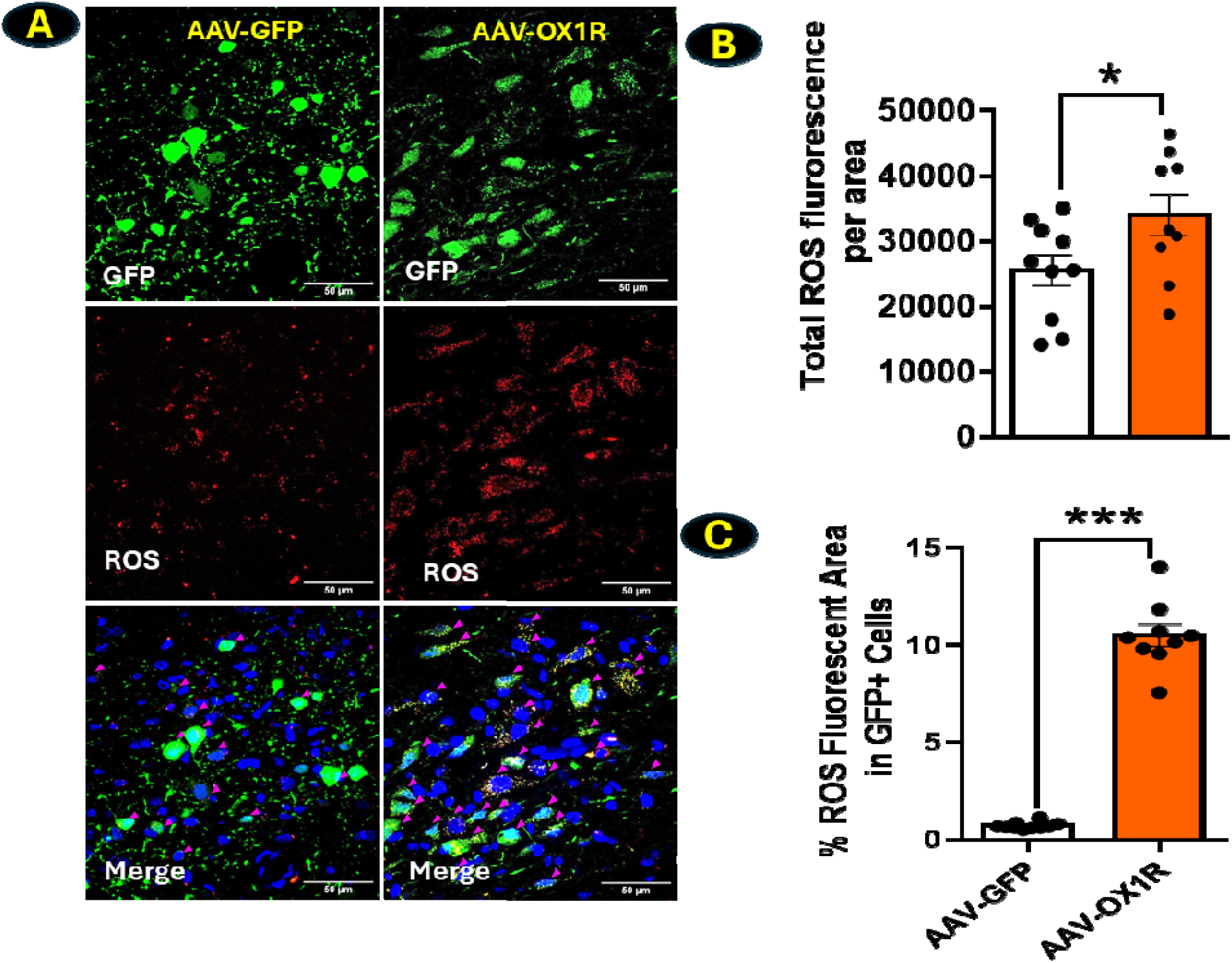
PVN OX1R overexpression increases ROS production in the PVN. MitoProbe was delivered via intracerebroventricular (ICV) injection to assess reactive oxygen species (ROS) production, followed by confocal imaging to quantify fluorescence in the PVN. **(A)** Representative images showing GFP expression and ROS signals in the PVN of AAV-OX1R and AAV-GFP rats. **(B)** Total ROS fluorescence was significantly elevated in AAV-OX1R rats compared with AAV-GFP controls (n = 5 per group, P < 0.05). **(C)** ROS co-localization within GFP⁺ cells—identifying OX1R-overexpressing neurons—was approximately ten-fold higher in the AAV-OX1R group (**P < 0.0001). Data are presented as mean ± SEM. Statistical comparisons were made using Student’s t-test. Scale bar: 50 µm.

### 4. PVN OX1R Overexpression Increases ROS Generation in the RVLM and Adrenal Gland

We further examined whether PVN OX1R overexpression affected two additional cardiovascular regulatory sites, the rostral ventrolateral medulla (RVLM) and adrenal gland, and assessed the co-localization of ROS (red) with OX1R-positive cells (green). Rats with OX1R overexpression showed a marked increase in red fluorescence in the RVLM (Figure 4A) and in both the adrenal cortex (Figure 4B1) and medulla (Figure 4B2), indicating elevated ROS levels. This was most apparent in the merged images, where strong yellow/orange signals reflected pronounced co-localization of OX1R and ROS (Figure 4). Together, these results suggest that PVN OX1R overexpression enhances oxidative stress not only within the PVN but also in other key cardiovascular regulatory regions.

**Figure 4.**
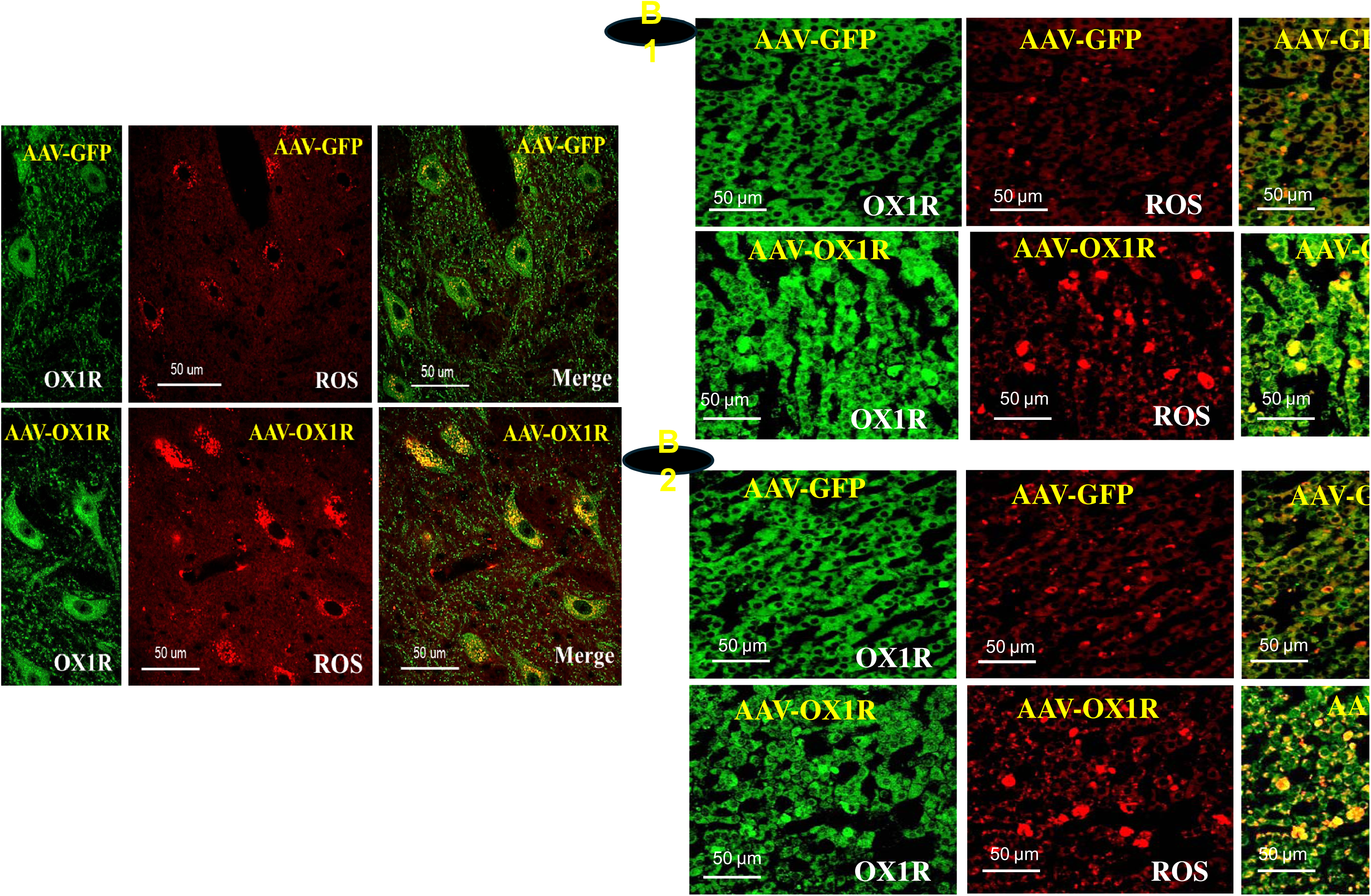
PVN OX1R overexpression elevates ROS levels in the RVLM and adrenal gland. **(A)** ROS fluorescence and its colocalization with OX1R in the RVLM. (**B**) ROS fluorescence and its colocalization with OX1R in the adrenal cortex (B1) and adrenal medulla (B2). In AAV2-GFP control rats, ROS fluorescence was low in both the RVLM and adrenal gland, with only minimal overlap with OX1R. In contrast, AAV2-OX1R rats displayed markedly increased ROS fluorescence and strong colocalization with OX1R (yellow/orange in merged images). n = 3 per group. Scale bar: 50 µm.

### 5. PVN OX1R Overexpression Significantly Increases Sympathetic Outflow and ABP in Response to Orexin A Administration

We further investigated the effect of OX1R activation on ABP and sympathetic outflow. Specifically, we examined whether orexin A administration into the PVN produces a greater increase in MAP and RSNA in rats with PVN OX1R overexpression compared with controls. Adult male SD rats were divided into two groups and received bilateral PVN injections of either AAV2-OX1R or control virus AAV2-GFP. Four to five weeks after injection, rats were anesthetized and received PVN microinjections of orexin A (30 pmol/150 nL/side). MAP and RSNA were recorded in each rat.

PVN administration of orexin A significantly increased MAP and RSNA in both OX1R-overexpressing and control rats; however, the magnitude of these responses was greater in OX1R-overexpressing rats (ΔRSNA: AAV-GFP, 72.7 ± 4.9%, n = 8 vs. AAV-OX1R, 155.4 ± 40.0%, n = 6; P < 0.05; ΔMAP: AAV-GFP, 16.3 ± 1.3 mmHg, n = 8 vs. AAV-OX1R, 26.6 ± 4.4 mmHg, n = 6; P < 0.05). No significant difference in HR was observed between groups (AAV-GFP, 10.3 ± 4.4 bpm, n = 8 vs. AAV-OX1R, 16.2 ± 6.2 bpm, n = 5; P > 0.05).

To determine whether the enhanced MAP and RSNA responses were specifically mediated by OX1R, OX1R-overexpressing rats were pretreated with the selective OX1R antagonist SB-408124 (30 nmol/side in 150 nL) before PVN orexin A administration. Pretreatment with SB-408124 markedly attenuated the orexin A–induced increase in RSNA (AAV2-OX1R, 155.4 ± 40.0%, n = 6 vs. SB-408124, 84.3 ± 17.6%, n = 5; P < 0.05) and MAP (AAV-OX1R, 26.6 ± 4.4 mmHg, n = 6 vs. SB408124, 12.4 ± 2.1 mmHg, P<0.05 (Figure 5).

**Figure 5.**
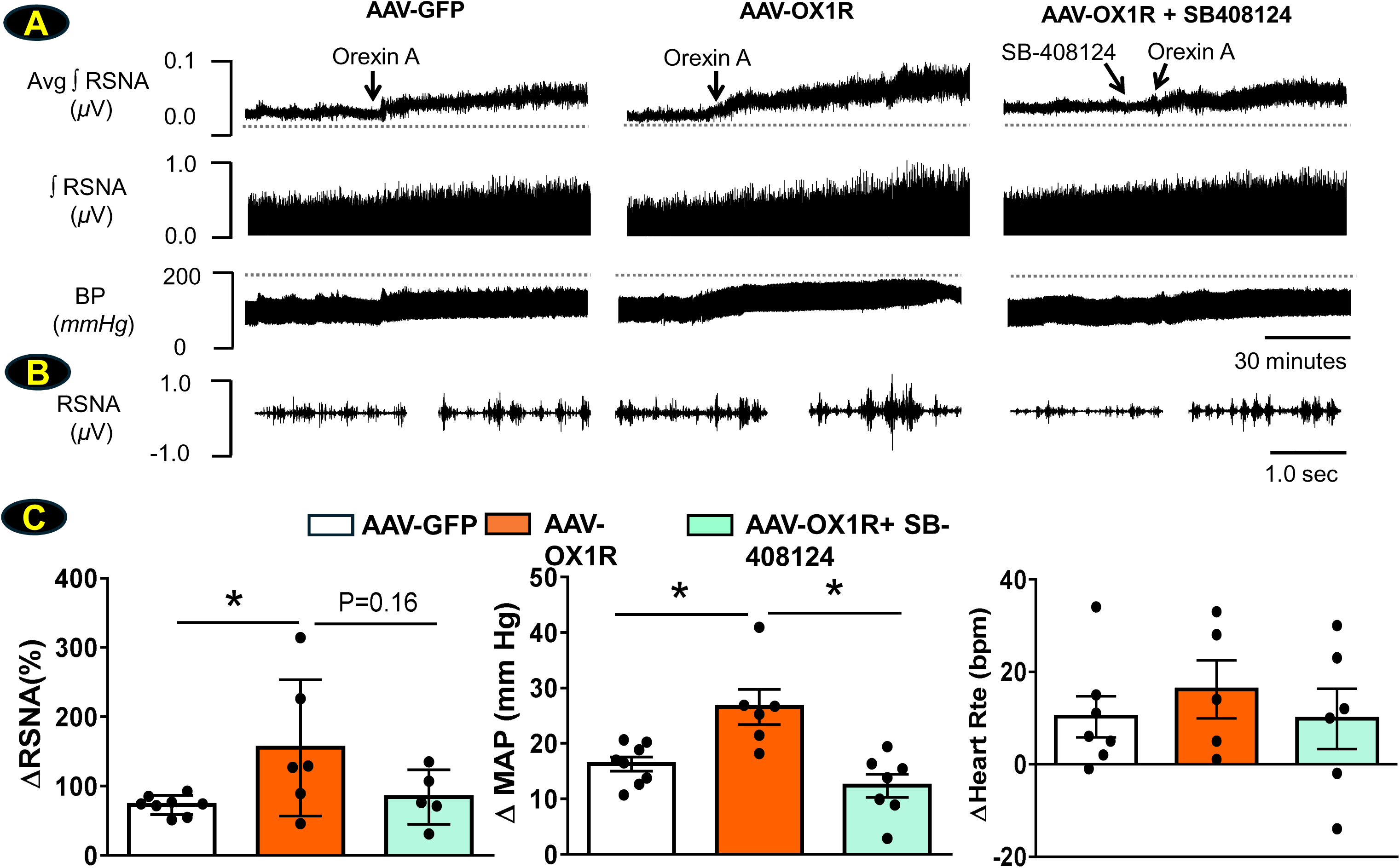
PVN OX1R Overexpression Enhances Sympathetic and Pressor Responses to Orexin. **A** Rats were randomly assigned to receive bilateral PVN injections of either AAV-OX1R or control virus. Five weeks post-injection, anesthetized rats underwent bilateral microinjections of orexin A (30 pmol/side) or SB-408124 (300 pmol) followed by orexin A (30 pmol) into the PVN. Mean arterial pressure (MAP) and renal sympathetic nerve activity (RSNA) were continuously recorded. (**A**) Representative traces showing RSNA and MAP responses to bilateral PVN microinjections of orexin A or SB-408124 followed by orexin A in AAV-GFP among AAV-OX1R rats. (**B**) Two-second sample traces of RSNA recorded before (left) and after (right) PVN injection of orexin A or SB-408124 followed by orexin A. **C**: Summary data showing changes in MAP, RSNA, and heart rate in response to PVN orexin A injection across experimental groups (AAV-GFP: n=8, AAV-OX1R; n=6, AAV-OX1R+SB-408124: n=5∼7, *P<0.05). Data are presented as mean ± SEM, and statistical analyses were performed using Student’s t-test.

### 6. PVN OX1R Overexpression Significantly Increases Plasma AVP Levels

Our previous work demonstrated that OX1R activation upregulates AVP expression in the PVN[5]. In the present study, we observed that PVN OX1R overexpression increased water intake and reduced urine output, suggesting that PVN OX1R overexpression alters AVP secretion. In addition, orexin A administration increased RSNA, indicating that PVN OX1R overexpression might also elevate plasma NE levels. To test these possibilities, we compared plasma AVP and NE levels between OX1R-overexpressing and control rats.

The result showed that plasma AVP was significantly elevated in the OX1R-overexpressing group compared with controls (AAV2-GFP: 4.4 ± 0.52 vs. AAV2-OX1R: 6.2 ± 0.3 pg/ml; n = 5; P < 0.05). However, there are no significant changes in plasma NE levels between PVN OX1R overexpression and control rats (AAV2-GFP: 4.6 ± 1.0 vs. AAV2-OX1R: 4.1± 1.1 ng/ml; n = 5; P > 0.05) (**Figure 6**).

**Figure 6.**
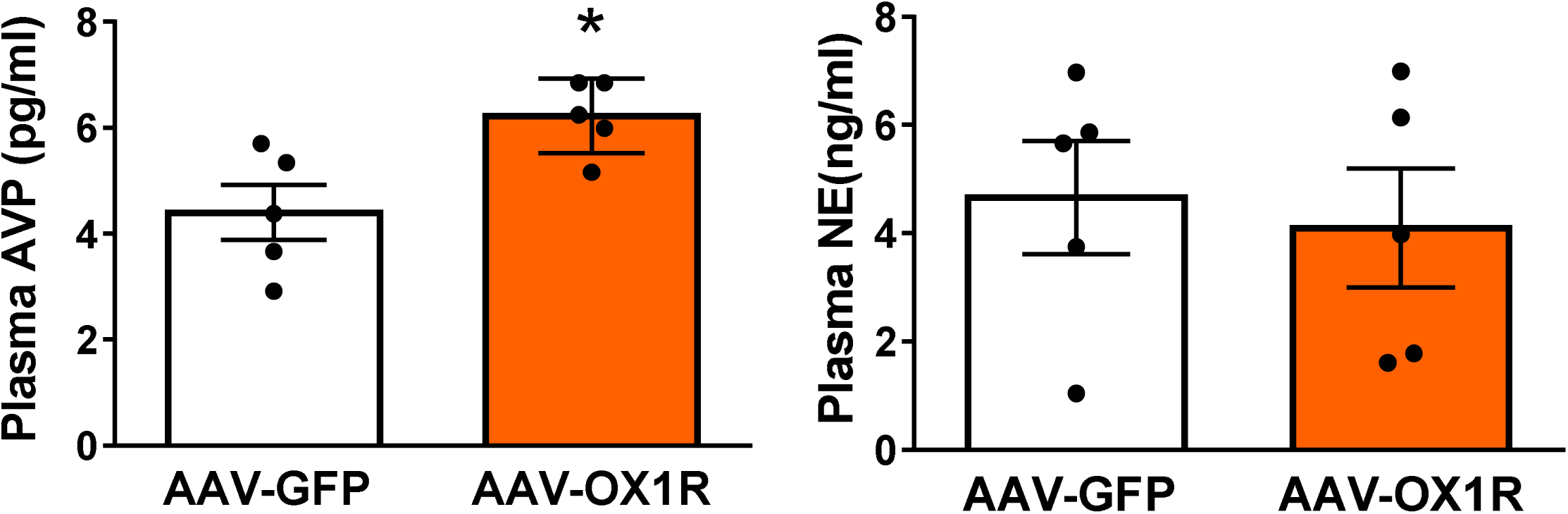
PVN OX1R overexpression increases plasma AVP levels. Compared with AAV2-GFP controls, PVN OX1R overexpression rats showed significantly higher plasma AVP levels (n=5 per group,* P < 0.05), with no significant changes in plasma norepinephrine (NE) levels (n=5 pergroup). Data are presented as mean ± SEM, and statistical analyses were performed using Student’s t-test.

### 7. AAV2-OX1R Increases Oxidative Stress and Excitability in Brain Neurons

Given that our in vivo PVN OX1R overexpression study demonstrated significantly increased ROS production localized to OX1R-positive neurons, and electrophysiological data showed elevated sympathetic outflow, we then investigated whether OX1R overexpression affects the expression of oxidative stress–related genes and neuronal excitation markers using brain neurons in primary culture. 3-day-old primary brain neuronal cultures from neonatal SD rats were infected with either AAV2-OX1R or control AAV2-GFP. Five days post-infection, cells were treated with orexin A or vehicle (saline). Six hours after treatment, neurons were collected for qPCR analysis of target gene expression. In OX1R-overexpressing neurons, SOD1 expression was significantly reduced, while other genes remain unchanged. Following orexin A stimulation, SOD2 expression was also significantly decreased. Moreover, orexin A treatment led to a significant upregulation of CYBB (1.7-fold), and neuronal excitation markers Jun (4-fold) and Fosl1 (7-fold) (Figure 7).

**Figure 7.**
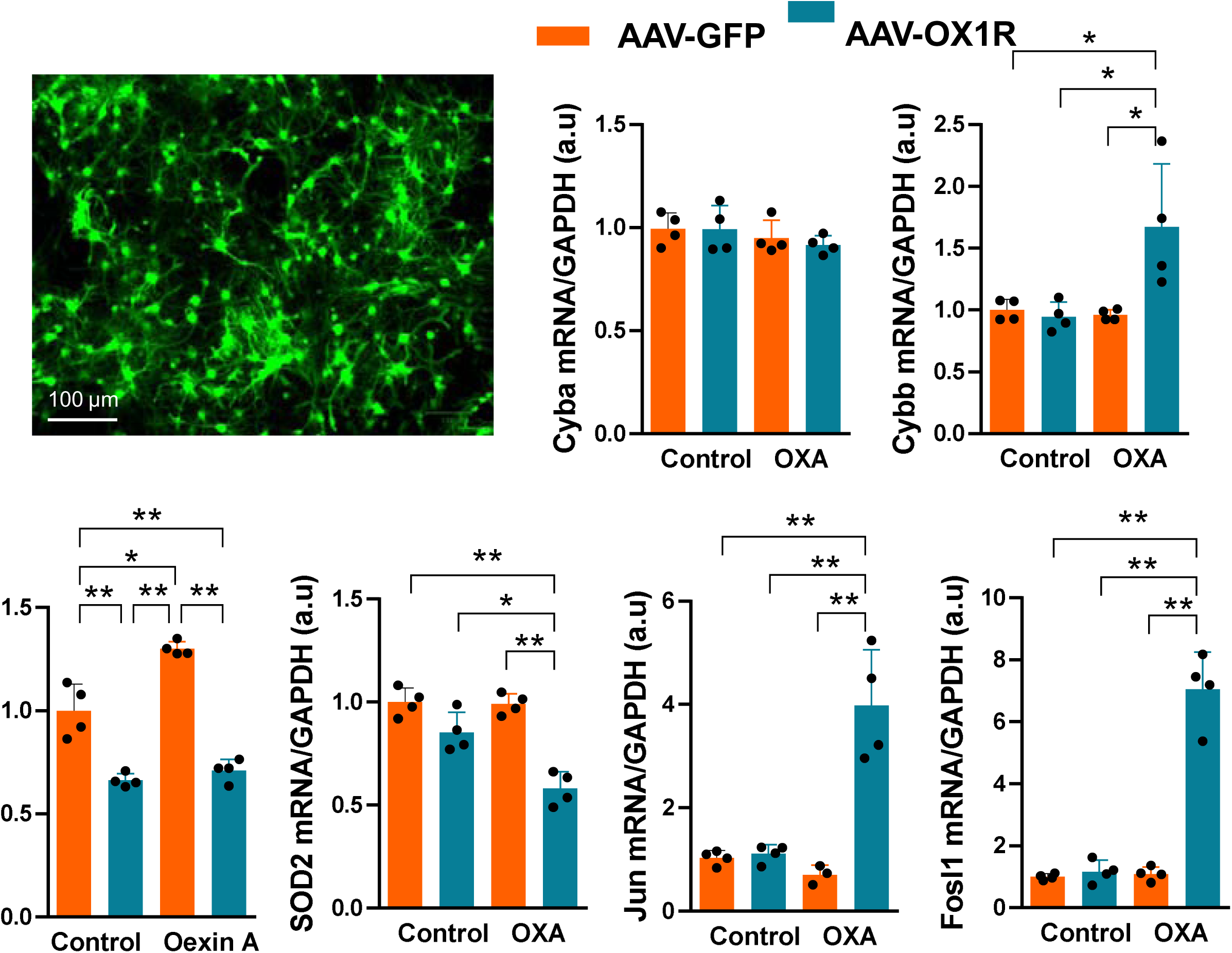
Effects of OX1R overexpression and orexin A stimulation on oxidative stress–related and neuronal excitation genes in primary neurons. Primary neuronal cultures derived from neonatal Sprague–Dawley rats were infected with either AAV2-OX1R or control virus (AAV2-GFP). Five days post-infection, cells were treated with orexin A or vehicle for 6 hours before quantitative PCR (qPCR) analysis. Fluorescent imaging confirmed reporter gene expression in AAV-OX1R–infected neurons, indicating successful viral transduction. Graphs summarize the effects of OX1R overexpression, with or without orexin A treatment, on the expression of Cyba, Cybb, SOD1, SOD2, and neuronal activation markers Jun and Fosl1. In OX1R-overexpressing neurons, basal SOD1 expression was reduced by ∼30% (*P<0.05), while other oxidative stress–related genes remained unchanged. Upon orexin A stimulation, SOD2 expression was significantly decreased by ∼35%, whereas CYBB (1.7-fold), Jun (4-fold), and Fosl1 (7-fold) were markedly upregulated (n=4 per group, *P<0.05) compared to control AAV-GFP infection neurons. Data are presented as mean ± SEM, and statistical analyses were performed using Student’s t-test. Scale bar: 100 µm.

## Discussion

The present study demonstrates that chronic overexpression of OX1R in the PVN of the hypothalamus contributes to sustained increases in arterial BP, elevated water intake, augmented sympathetic outflow, enhanced oxidative stress, and increased plasma AVP level. Together, these findings provide new mechanistic insights into the role of central orexin signaling in cardiovascular regulation and the development of hypertension.

### PVN OX1R Overexpression and Cardiovascular Regulation

Our data show that PVN-targeted OX1R overexpression induced a mild but persistent elevation in ABP that became apparent within two weeks and was maintained throughout the study period. Heart rate remained largely unaffected. This observation suggest that chronic overexpression of OX1R in the PVN of normal rat may play a crucial role in modulating cardiovascular tone without necessarily altering basal heart rate. The concurrent increase in water intake, together with elevated plasma AVP levels, indicates that PVN OX1R signaling also affects fluid balance and neuroendocrine regulation, reinforcing the multifaceted role of this receptor in homeostatic control.

### Oxidative Stress as a Mechanistic Link

A major finding of this study is that OX1R overexpression markedly increased reactive oxygen species (ROS) production within PVN neurons. Confocal imaging revealed pronounced ROS accumulation specifically in OX1R-expressing cells, implicating oxidative stress as a central mediator of the hypertensive phenotype. These results are consistent with prior reports showing that central oxidative stress contributes to neurogenic hypertension[8, 15, 16]. Supporting this, gene expression analysis revealed a redox imbalance characterized by decreased SOD1 and increased Cybb, further indicating a shift toward a pro-oxidant state. Notably, enhanced ROS production was not confined to the PVN but was also evident in the rostral ventrolateral medulla (RVLM) and adrenal gland—key regions involved in sympathetic regulation and control of cardiovascular function.

The precise mechanism by which OX1R overexpression drives oxidative stress beyond the PVN remains unclear. However, it is well established that the PVN regulates SNA through presympathetic neurons projecting either to the RVLM or directly to the intermediolateral column (IML) of the spinal cord. Elevated ROS within the PVN may enhance excitatory signaling along these pathways, thereby increasing neuronal excitability and ROS generation within the RVLM. In turn, this increased sympathetic output could excessively activate the adrenal gland, further augmenting oxidative stress in peripheral tissues. Collectively, these findings suggest that oxidative stress originating in the PVN initiates a feed-forward cascade, extending redox imbalance from hypothalamic circuits to brainstem autonomic nucleus and ultimately to endocrine targets. This network-wide propagation of oxidative stress highlights PVN OX1R signaling as a potential amplifier of sympathoexcitation and cardiovascular dysfunction

### Sympathetic Outflow and Responsiveness to Orexin A

To evaluate the impact of OX1R overexpression in the PVN on sympathetic outflow, we microinjected the OX1R agonist orexin A into the PVN of OX1R-overexpressing rats and control AAV-GFP rats. ABP and RSNA were recorded, and the results showed that orexin A administration increased ABP and RSNA in both groups. However, this response was markedly exaggerated in the PVN OX1R-overexpressing rats. Importantly, pretreatment with the selective OX1R antagonist SB-408124 attenuated these exaggerated responses in PVN OX1R overexpression rats, confirming receptor-specific mediation. Prior studies have demonstrated that orexin A in the PVN and RVLM promotes sympathoexcitation and elevates arterial pressure[5, 17]. The present findings extend these observations by showing that OX1R overexpression sensitizes PVN neurons to orexin A, thereby amplifying sympathoexcitatory signaling and contributing to sustained hypertension.

### Hormonal Contributions to ABP Elevation

Our results also demonstrated that plasma AVP levels were significantly elevated in OX1R-overexpressing rats compared with control rats. This finding suggests that neuroendocrine mechanisms contribute to the observed increase in ABP. Although still debated, AVP is widely recognized as a potent vasoconstrictor and antidiuretic hormone. Consistent with this role, OX1R-overexpressing rats exhibited increased water intake and reduced urine output under water deprivation, both of which align with elevated plasma AVP levels. Notably, elevated AVP concentrations have been implicated in several models of experimental hypertension [6, 18]. Therefore, OX1R overexpression in the PVN appears to disrupt not only autonomic regulation but also neuroendocrine homeostasis, producing a synergistic effect on ABP control.

Interestingly, we did not observe a corresponding increase in plasma NE levels, even though PVN administration of orexin A markedly increased renal sympathetic nerve activity (RSNA) in OX1R-overexpressing rats compared with controls (Figure 5). One possible explanation for this discrepancy is that NE spillover into the circulation may be counterbalanced by enhanced neuronal reuptake or metabolism[19, 20]. Alternatively, the sympathetic activation induced by PVN OX1R overexpression may be regionally localized, which primarily affects renal sympathetic nerves, thereby minimizing changes in systemic plasma NE concentrations.

### Cellular Mechanisms in Primary Neurons

Our in vivo experiments revealed a robust increase in ROS production within the PVN of OX1R-overexpressing rats, with ROS primarily colocalized to OX1R-positive neurons. This suggests that OX1R overexpression directly promotes oxidative stress in these cells. Interestingly, gene expression assays did not show statistically significant changes in major ROS-generating or antioxidant enzymes such as NADPH oxidase and SOD, although trends toward increased Cybb and decreased SOD1 were observed. Given the cellular heterogeneity of the PVN—including astrocytes, microglia, and neurons—and the possibility of contamination from surrounding tissue during punch sampling, we employed primary neuronal cultures to more specifically examine OX1R-dependent effects.

Our in vitro findings reinforced the in vivo results by showing that OX1R overexpression in neurons enhances both oxidative stress and excitability. In OX1R overexpression neurons, following orexin A stimulation, expression of the mitochondrial specific antioxidant enzyme gene SOD2[21] was suppressed, whereas Cybb, a subunit of NADPH oxidase and a major contributor to ROS generation, was markedly upregulated. In parallel, the neuronal activation marker Jun was strongly induced, consistent with increased oxidative burden and heightened neuronal activity. These observations agree with previous work demonstrating that orexin signaling promotes excitability through calcium influx and activation of intracellular signaling cascades[22, 23]. Collectively, the oxidative and excitatory changes provide a mechanistic framework for the enhanced sympathetic outflow and elevated ABP observed in vivo.

## Conclusion

In summary, chronic PVN OX1R overexpression drives sustained hypertension through a multifactorial mechanism involving increased sympathetic outflow, heightened oxidative stress, and altered neuroendocrine signaling. These findings identify OX1R as a critical modulator of autonomic and hormonal regulation within the PVN and suggest that central orexin signaling represents a novel therapeutic target for the treatment of neurogenic hypertension.

## Translational Perspective

This study provides critical insight into the central mechanisms linking orexin system hyperactivity to hypertension. By demonstrating that chronic OX1R overexpression in the PVN elevates blood pressure, sympathetic nerve activity, vasopressin release, and oxidative stress, these findings reveal a potential neurogenic pathway contributing to sustained hypertension. Clinically, targeting OX1R signaling may offer a novel therapeutic strategy for patients with resistant hypertension. Modulation of orexin pathways could also help mitigate associated metabolic and neuroendocrine dysregulation, offering a more comprehensive approach to cardiovascular disease prevention and treatment rooted in central nervous system mechanisms.

## Funding Support

This work was supported by the National Institute of Health [R01HL163159 to Z.S., R15EB035866 to L.B., and R15HL177713 to R.L].

## Conflict of Interest

All authors declare that there are no conflicts of interest regarding publication of this work.

## Notes

### Competing Interest Statement

The authors have declared no competing interest.

## References

[1] L.W. Swanson, P.E. Sawchenko, Paraventricular nucleus: a site for the integration of neuroendocrine and autonomic mechanisms, Neuroendocrinology 31(6) (1980) 410–7.

[2] E.E. Benarroch, Paraventricular nucleus, stress response, and cardiovascular disease, Clin Auton Res 15(4) (2005) 254–63.

[3] J.J. Zhou, F. Yuan, Y. Zhang, D.P. Li, Upregulation of orexin receptor in paraventricular nucleus promotes sympathetic outflow in obese Zucker rats, Neuropharmacology 99 (2015) 481–90.

[4] T. Sakurai, A. Amemiya, M. Ishii, I. Matsuzaki, R.M. Chemelli, H. Tanaka, S.C. Williams, J.A. Richarson, G.P. Kozlowski, S. Wilson, J.R. Arch, R.E. Buckingham, A.C. Haynes, S.A. Carr, R.S. Annan, D.E. McNulty, W.S. Liu, J.A. Terrett, N.A. Elshourbagy, D.J. Bergsma, M. Yanagisawa, Orexins and orexin receptors: a family of hypothalamic neuropeptides and G protein-coupled receptors that regulate feeding behavior, Cell 92(5) (1998) 1 page following 696.

[5] M.J. Huber, Y. Fan, E. Jiang, F. Zhu, R.A. Larson, J. Yan, N. Li, Q.H. Chen, Z. Shan, Increased activity of the orexin system in the paraventricular nucleus contributes to salt-sensitive hypertension, Am J Physiol Heart Circ Physiol 313(6) (2017) H1075–H1086.

[6] J.A. Bigalke, H. Gao, Q.H. Chen, Z. Shan, Activation of Orexin 1 Receptors in the Paraventricular Nucleus Contributes to the Development of Deoxycorticosterone Acetate-Salt Hypertension Through Regulation of Vasopressin, Front Physiol 12 (2021) 641331.

[7] Y. Fan, E. Jiang, T. Hahka, Q.H. Chen, J. Yan, Z. Shan, Orexin A increases sympathetic nerve activity through promoting expression of proinflammatory cytokines in Sprague Dawley rats, Acta Physiol (Oxf) 222(2) (2018).

[8] T. Kishi, Y. Hirooka, Oxidative stress in the brain causes hypertension via sympathoexcitation, Front Physiol 3 (2012) 335.

[9] S. Chopra, C. Baby, J.J. Jacob, Neuro-endocrine regulation of blood pressure, Indian J Endocrinol Metab 15 Suppl 4(Suppl4) (2011) S281–8.

[10] T. Kishi, Y. Hirooka, Y. Kimura, K. Ito, H. Shimokawa, A. Takeshita, Increased reactive oxygen species in rostral ventrolateral medulla contribute to neural mechanisms of hypertension in stroke-prone spontaneously hypertensive rats, Circulation 109(19) (2004) 2357–62.

[11] J. Chen, C. Xia, J. Wang, M. Jiang, H. Zhang, C. Zhang, M. Zhu, L. Shen, D. Zhu, The effect of orexin-A on cardiac dysfunction mediated by NADPH oxidase-derived superoxide anion in ventrolateral medulla, PLoS One 8(7) (2013) e69840.

[12] S. Kowalewski, K. Czarzasta, L. Puchalska, E. Szczepanska-Sadowska, A. Wsol, A. Cudnoch-Jedrzejewska, Interaction of Orexin A and Vasopressin in the Brain Plays a Role in Blood Pressure Regulation in WKY and SHR Rats, Med Sci Monit 26 (2020) e926825.

[13] R.L. Thunhorst, B. Xue, T.G. Beltz, A.K. Johnson, Age-related changes in thirst, salt appetite, and arterial blood pressure in response to aldosterone-dexamethasone combination in rats, Am J Physiol Regul Integr Comp Physiol 308(10) (2015) R807–15.

[14] H. Gao, J. Bigalke, E. Jiang, Y. Fan, B. Chen, Q.H. Chen, Z. Shan, TNFalpha Triggers an Augmented Inflammatory Response in Brain Neurons from Dahl Salt-Sensitive Rats Compared with Normal Sprague Dawley Rats, Cell Mol Neurobiol 42(6) (2022) 1787–1800.

[15] A. Nagae, M. Fujita, H. Kawarazaki, H. Matsui, K. Ando, T. Fujita, Sympathoexcitation by oxidative stress in the brain mediates arterial pressure elevation in obesity-induced hypertension, Circulation 119(7) (2009) 978–86.

[16] M. Fujita, K. Ando, A. Nagae, T. Fujita, Sympathoexcitation by oxidative stress in the brain mediates arterial pressure elevation in salt-sensitive hypertension, Hypertension 50(2) (2007) 360–7.

[17] I.Z. Shahid, A.A. Rahman, P.M. Pilowsky, Orexin A in rat rostral ventrolateral medulla is pressor, sympatho-excitatory, increases barosensitivity and attenuates the somato-sympathetic reflex, Br J Pharmacol 165(7) (2012) 2292–303.

[18] M. Yu, V. Gopalakrishnan, J.R. McNeill, Role of endothelin and vasopressin in DOCA-salt hypertension, Br J Pharmacol 132(7) (2001) 1447–54.

[19] M. Esler, G. Jennings, P. Korner, P. Blombery, N. Sacharias, P. Leonard, Measurement of total and organ-specific norepinephrine kinetics in humans, Am J Physiol 247(1 Pt 1) (1984) E21–8.

[20] Y. Sata, G.A. Head, K. Denton, C.N. May, M.P. Schlaich, Role of the Sympathetic Nervous System and Its Modulation in Renal Hypertension, Front Med (Lausanne) 5 (2018) 82.

[21] A.M. Dosunmu-Ogunbi, K.C. Wood, E.M. Novelli, A.C. Straub, Decoding the role of SOD2 in sickle cell disease, Blood Adv 3(17) (2019) 2679–2687.

[22] J.X. Xia, S.Y. Fan, J. Yan, F. Chen, Y. Li, Z.P. Yu, Z.A. Hu, Orexin A-induced extracellular calcium influx in prefrontal cortex neurons involves L-type calcium channels, J Physiol Biochem 65(2) (2009) 125–36.

[23] Y. Fan, E. Jiang, H. Gao, J. Bigalke, B. Chen, C. Yu, Q. Chen, Z. Shan, Activation of Orexin System Stimulates CaMKII Expression, Front Physiol 12 (2021) 698185.

